# Inhibition of CMP-sialic acid transport by endogenous 5-methyl CMP

**DOI:** 10.1101/2020.06.30.180356

**Authors:** Shivani Ahuja, James Cahill, Kimberly Hartfield, Matthew R. Whorton

## Abstract

Nucleotide-sugar transporters (NSTs) transport nucleotide-sugar conjugates into the Golgi lumen where they are then used in the synthesis of glycans. We previously reported crystal structures of a mammalian NST, the CMP-sialic acid transporter (CST) (Ahuja and Whorton 2019). These structures elucidated many aspects of substrate recognition, selectivity, and transport; however, one fundamental unaddressed question is how the transport activity of NSTs might be physiologically regulated as a means to produce the vast diversity of observed glycan structures. Here, we describe the discovery that an endogenous methylated form of cytidine monophosphate (m^5^CMP) binds and inhibits CST. The presence of m^5^CMP in cells results from the degradation of RNA that has had its cytosine bases post-transcriptionally methylated through epigenetic processes. Therefore, this work not only demonstrates that m^5^CMP represents a novel physiological regulator of CST, but it also establishes a link between epigenetic control of gene expression and regulation of glycosylation.

## Introduction

Glycosylation is the most common form of protein and lipid modification (Dwek, Butters et al. 2002, Ohtsubo and Marth 2006, Stanley 2011, Moremen, Tiemeyer et al. 2012). Glycosylation affects protein folding, stability, and activity, and therefore impacts nearly every aspect of biology. Most glycosylation occurs in the ER and Golgi lumens, where glycosyltransferase enzymes build glycan chains by transferring sugars from nucleotide-coupled sugar donors to glycan acceptors. For these reactions to occur, nucleotide-coupled sugars must be transported from the cytoplasm, where they are synthesized, across the ER and Golgi membranes and into the lumenal space. This is accomplished by a family of proteins called nucleotide sugar transporters (NSTs) (Ishida and Kawakita 2004, Song 2013, Hadley, Maggioni et al. 2014).

Glycan structures can be very complex and also highly diverse (Moremen, Tiemeyer et al. 2012). A single type of protein may be differentially glycosylated across different cell types or even within the same cell, which can result in differing functional effects. Many factors are thought to affect the generation of diverse glycosylation patterns, with one being the availability of nucleotide sugars in the Golgi lumen. One of the primary factors that controls the availability of nucleotide sugars in the Golgi lumen is the transport activity of NSTs, yet little is known about what cellular processes may regulate NST activity.

It has been shown that free nucleotide monophosphates (NMPs) can inhibit uptake of nucleotide sugars into the Golgi lumen, ostensibly by competing with the nucleotide sugar for binding to the NST (Chiaramonte, Koviach et al. 2001). While it is known that concentrations of NMPs with canonical bases (A, T, C, G, or U) can vary between cell type and fluctuate depending on the metabolic needs of a cell (Traut 1994), there are currently no established links between these fluctuations and regulation of glycosylation. However, it has recently been appreciated that pools of cellular NMPs are comprised of more than just the canonical bases (Zeng, Qi et al. 2017). Enzymatic degradation of DNA and RNA that has been modified as part of epigenetic control of gene expression can contribute a vast diversity to cellular pools of free NMPs, since there are hundreds of known ways that DNA and RNA bases may be modified (Liu and Pan 2015, Chen, Zhao et al. 2016, Zhao, Roundtree et al. 2016). One of the most common ways that DNA and RNA are modified is through methylation of cytosine bases. Methylated cytosine in DNA has many well established physiological roles in controlling gene transcription (Chen, Zhao et al. 2016), and although not as well understood, methylation of cytosine bases in various types of RNA is also very abundant (Squires, Patel et al. 2012, Gkatza, Castro et al. 2019, Trixl and Lusser 2019).

Cytosine is methylated at the C-5 position of the pyrimidine ring by a family of methyltransferases (Goll and Bestor 2005). It appears that these enzymes only recognize and methylate cytosine bases within the context of RNA or DNA polymers, rather than acting on free nucleotides. This means that the cellular concentration of free 5-methyl CMP (m^5^CMP) or 5-methyl deoxy-CMP (m^5^dCMP) principally originates from the breakdown of methylated RNA or DNA, respectively. Therefore, levels of free m^5^CMP or m^5^dCMP are a result of a combination of factors that control the level of RNA/DNA methylation, but also factors that control RNA/DNA decay.

The CMP-sialic acid transporter (CST) is one of seven types of mammalian NSTs and is highly selective for transporting CMP-sialic acid (CMP-Sia) into the Golgi lumen as well as for transporting the free nucleotide byproduct of the glycosyltransferase reaction, CMP, back to the cytoplasm. Sialic acid is commonly the terminal sugar added to glycan chains and has many functional roles (Varki 2008, Varki and Schauer 2009). Here, we describe the discovery of a molecule that co-purifies with CST. We use analytical HPLC, UV-Vis spectroscopy, and LC-MS/MS to identify this molecule as m^5^CMP. We show that m^5^CMP inhibits CMP-Sia transport, and a crystal structure of CST in complex with m^5^CMP provides insight into the molecular mechanism of high-affinity interaction between m^5^CMP and CST. Considering that m^5^CMP inhibits CMP-Sia transport and that m^5^CMP cellular concentrations are primarily related to post-transcriptional methylation of RNA, these results suggest a link between RNA epigenetics and regulation of cellular glycosylation.

## Results

### Initial characterization of a molecule that co-purifies with CST

After determining the structures of CST in complex with its two primary physiological substrates (CMP-Sia and CMP) (Ahuja and Whorton 2019), one of our next aims was to determine the structure of CST in the absence of any ligand. Our hope was that such a structure would help elucidate some of the conformational transitions that occur within CST upon ligand binding. Our approach to determine the structure of a ligand-free CST was to simply crystallize purified CST without the addition of any ligand. We were able to grow crystals of CST under these conditions. They were very small but they still allowed us to collect a partially-complete dataset with a resolution of 3.3 Å. Unexpectedly, the molecular replacement solution (data not shown) indicated that either CMP or a CMP-like molecule was still bound in the substrate-binding cavity.

We hypothesized that one reason a CMP-like molecule would be present in these crystals is if it was co-purified with CST. To test this hypothesis, we performed a phenol-chloroform extraction on a sample of purified CST to precipitate the protein and liberate any bound molecule. The aqueous fraction of this extract was run on a C18 HPLC column (Figure 1A). We observed a peak with a retention time of ~3.5 min, which was significantly different than the retention times of CMP, UMP, or CMP-Sia – a selection of candidate co-purifying molecules.

**Figure 1.**
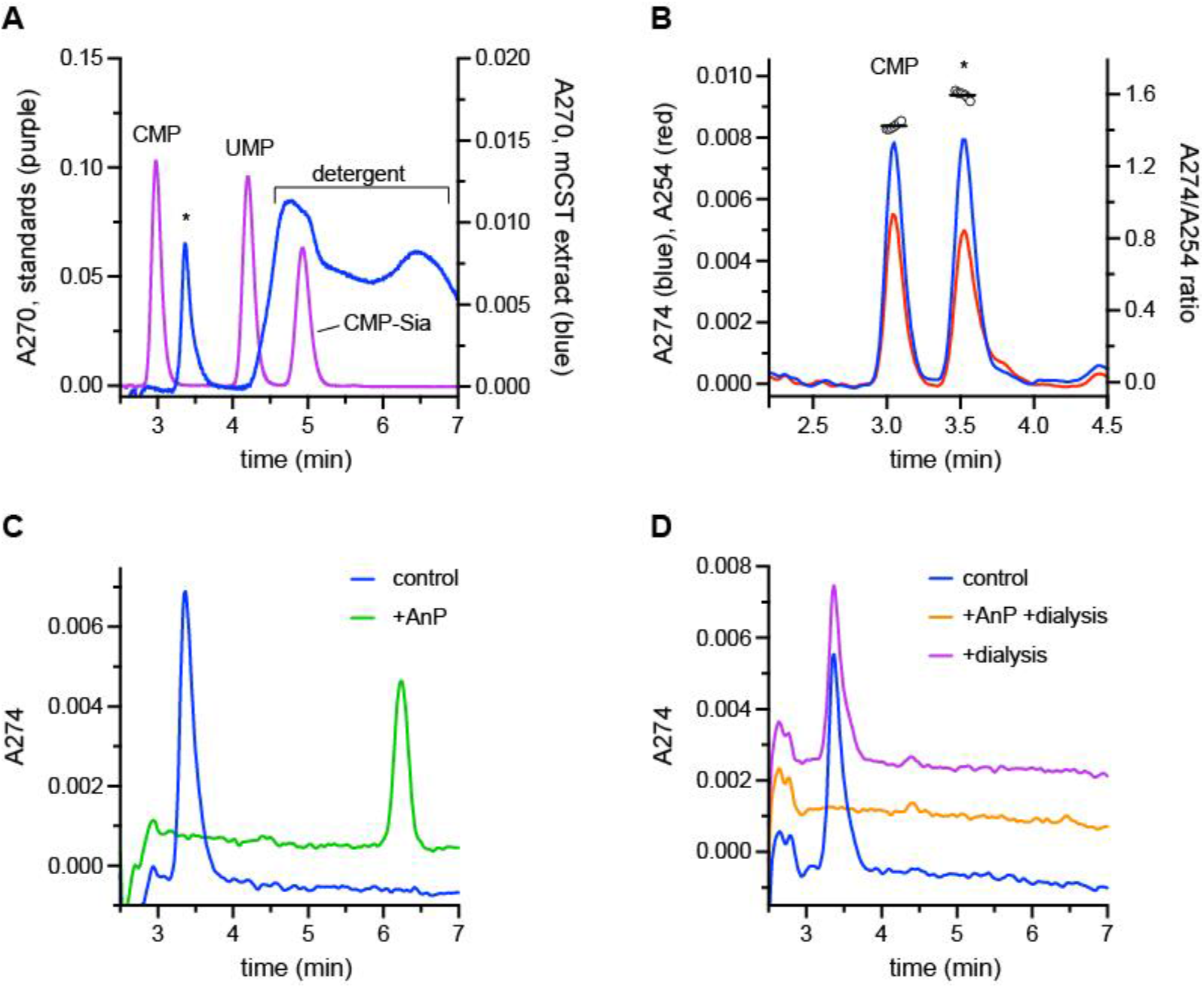
HPLC characterization of the molecule that co-purifies with CST. **A)** The aqueous layer of a phenol-chloroform extraction of purified CST was run on a C18 HPLC column (blue trace). The peak representing the unknown molecule is labeled with an ‘*’ and the broad peak representing residual detergent left in the extract is also labeled. A separate sample containing 50 μM each of CMP, UMP, and CMP-Sia was also run (purple trace). Peaks corresponding to each standard are labeled. **B)** A sample of a phenol-chloroform extract of CST with an extra 2.5 μM CMP added was run on a HPLC C18 column. Absorbance at 274 nm (blue trace) and 254 nm (red trace) was monitored. The ratio of the absorbance at these two wavelengths for a given time is plotted as the open circles on the right y-axis. The black horizontal lines represent the average A274/A254 ratio for the two peaks. **C)** A sample of a phenol-chloroform extract of CST was run on a C18 HPLC column after it had been incubated for 24 hr at 4°C either with (green trace) or without AnP (blue trace). **D)** A phenol-chloroform extraction was performed on a CST sample immediately after purification (blue trace), after it had undergone 24 hr dialysis (purple trace), or after it had undergone 24 hr dialysis in the presence of AnP (orange trace). The extracts were then run on C18 HPLC column. The traces in panels C and D are arbitrarily separated by either 0.001 (panel C) or 0.002 (panel D) units along the y-axis to improve clarity.

To further characterize this unknown peak, we next took a sample of the CST phenol-chloroform extract and added 2.5 μM CMP. This sample was run on the C18 HPLC column while we monitored the absorbance at both 274 nm and 254 nm (Figure 1B). Like the CMP peak, the peak at 3.5 min had a higher absorbance at 274 nm compared to 254 nm, indicating that the molecule has an aromatic group. Calculation of the A274/A254 ratio across the two peaks further shows that the peak at 3.5 min represents a different molecule than CMP since its A274/A254 ratio is 1.60 ± 0.02 compared to 1.42 ± 0.02 for CMP.

Considering that the unknown molecule has an aromatic group and has a similar retention time as CMP, we also wanted to see if it had a phosphate group. To do so, we treated the CST phenol-chloroform extract with a non-selective nucleotide phosphatase, Antarctic phosphatase (AnP). As seen in Figure 1C, AnP-treatment yielded a peak with a retention time of ~6.25 min. This shifted retention time indicates that a molecule with a unique chemical composition was formed, which is consistent with the generation of a new phosphate-lacking molecule. We also found that we still observe the peak at ~3.5 min even after a sample of purified CST has been dialyzed for 24 hr (Figure 1D) before being subjected to phenol-chloroform extraction. This is consistent with this unknown molecule having a high binding affinity for CST and explains how it stays bound over the course of a two-day purification procedure. However, if we include AnP during the dialysis, we no longer observe a peak at either 3.5 min or 6.25 min (Figure 1D). This indicates that the loss of the phosphate from the unknown molecule significantly reduces its binding affinity to the point where it can be dialyzed away. This is similar to our previous observation of how the removal of phosphate from CMP to form cytidine completely eliminates its ability to bind CST (Ahuja and Whorton 2019).

### Identification of the unknown molecule as 5-methyl CMP

To gain further insight into the identity of the molecule, we sent samples to the Northwest Metabolomics Research Center at the University of Washington to be analyzed by HPLC coupled to mass spectrometry (LC-MS), using electrospray ionization (ESI). A comparison of total ion current chromatograms from phenol-chloroform extracts of a buffer-only sample versus a sample containing purified CST protein did not reveal any peaks that were obviously-unique to the protein-containing sample (Figure 2 – figure supplement 1). Several peaks that were unique to the buffer-only control were observed; however, this may be partially due to the high concentration of denatured protein in the CST sample affecting the partitioning of some buffer components during the phenol-chloroform extraction.

An analysis of extracted ion chromatograms with a *m/z* range of 338.0701 ± 0.5 revealed a peak that was found only in the protein-containing sample (Figure 2A). Mass spectra that correspond to the retention time of this peak (10.7-10.8 min) are shown for the buffer-only sample (Figure 2B) and the protein-containing sample (Figure 2C). These clearly show that there is an ion with an *m/z* of 338.0722 (positive ESI mode) that is unique to the protein-containing sample. Similar analyses performed in negative ESI mode showed a unique ion with an *m/z* of 336.0631 (Figure 2D). Although there was a unique peak in the buffer-only sample in the extracted ion chromatogram (Figure 2A), the mass spectrum corresponding to the retention time for this peak (9.6-9.8 min) showed that the contributing ion has an *m/z* of 338.0457 (Figure 2E). This likely represents one of several ions that are unique to the buffer-only sample, as described above.

**Figure 2.**
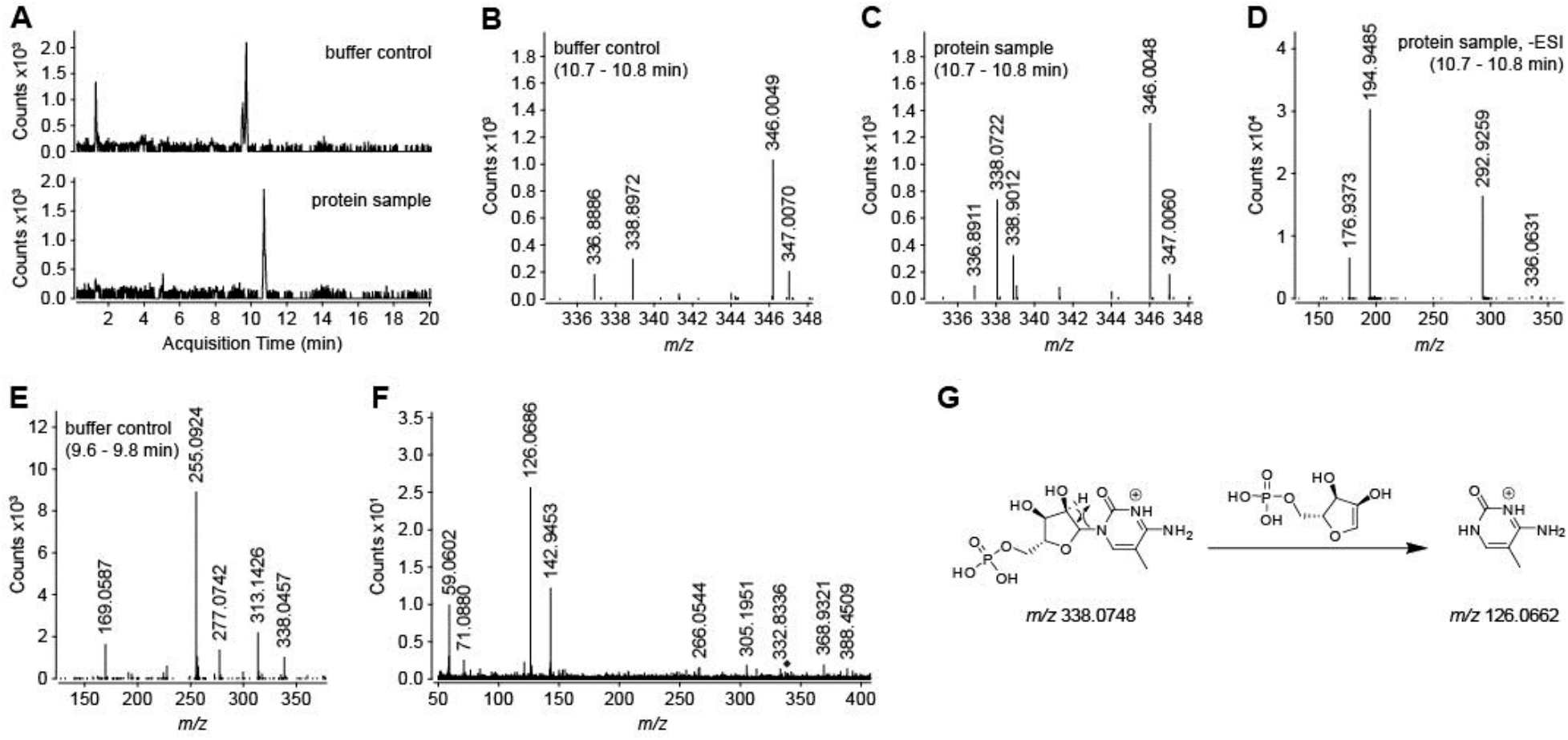
Comparison of mass spectra for buffer-only versus CST protein-containing samples. **A)** Extracted ion chromatograms for an *m/z* of 338.0701 ± 0.5 were taken from the LC-MS runs in positive ESI mode (shown in Figure 2 – figure supplement 1) and are shown for the buffer-only control (top) or the CST protein-containing sample (bottom). **B-E)** Mass spectra in either positive (panels B, C, and E) or negative (panel D) ESI modes are shown. Panels B-D show the spectra for the sample eluting from the HPLC column from between 10.7-10.8 min for the buffer-only control (B) or protein-containing samples (C and D). Panel E shows the spectrum for the buffer-only sample eluting from the HPLC column between 9.6-9.8 min. **F)** Tandem MS/MS was performed on the ion with *m/z* of 338.0722 from panel C, with a collision energy of 10 eV. A selection filter of 1.3 *m/z* was used. The black diamond indicates the position of the precursor ion. **G)** A putative scheme describes the fragmentation of m^5^CMP ([M+H]^+^ = 338.0748) to produce the most prominent ion (m/z of 126.0686) in the MS/MS spectrum shown in panel F.

To further characterize this ion that was unique to the protein-containing sample, it was subjected to MS/MS fragmentation in positive ESI mode (Figure 2F). The most abundant product ion had an *m/z* of 126.0686, which supported an annotation of the precursor ion as 5-methyl-cytidine 5’-monophosphate (m^5^CMP), according to the scheme shown in Figure 2G. m^5^CMP has a monoisotopic mass of 337.0675 Da which equates to an *m/z* of 338.0748 and 336.0602 in positive and negative ESI modes, respectively. This gives ppm errors of 7.6 and 8.6, respectively, with regards to the measured *m/z*’s stated above, which is within the 5-10 ppm mass accuracy of the MS instrument that was used (Bristow and Webb 2003). The annotation of the precursor ion as m^5^CMP does not account for the other prominent ions in the MS/MS spectrum (Figure 2F); however, it is possible that they derive from the ion with an *m/z* of 338.9012 (Figure 2C), which would have been included in the MS/MS fragmentation since the precursor ion selection filter had an *m/z* isolation width of 1.3.

To reconcile this finding with our HPLC observations, we prepared a sample containing 5 μM each of CMP and m^5^CMP. We ran this sample as well as a phenol-chloroform extract of CST on a C18 HPLC column (Figure 3A). We saw that m^5^CMP eluted with a nearly identical retention time as that of the molecule that co-purifies with CST. We also characterized the A274/A254 ratio for m^5^CMP and found it to be 1.60 ± 0.01 (Figure 3B), which is essentially identical to the A274/A254 ratio measured for the molecule that co-purifies with CST (Fig. 1B).

**Figure 3.**
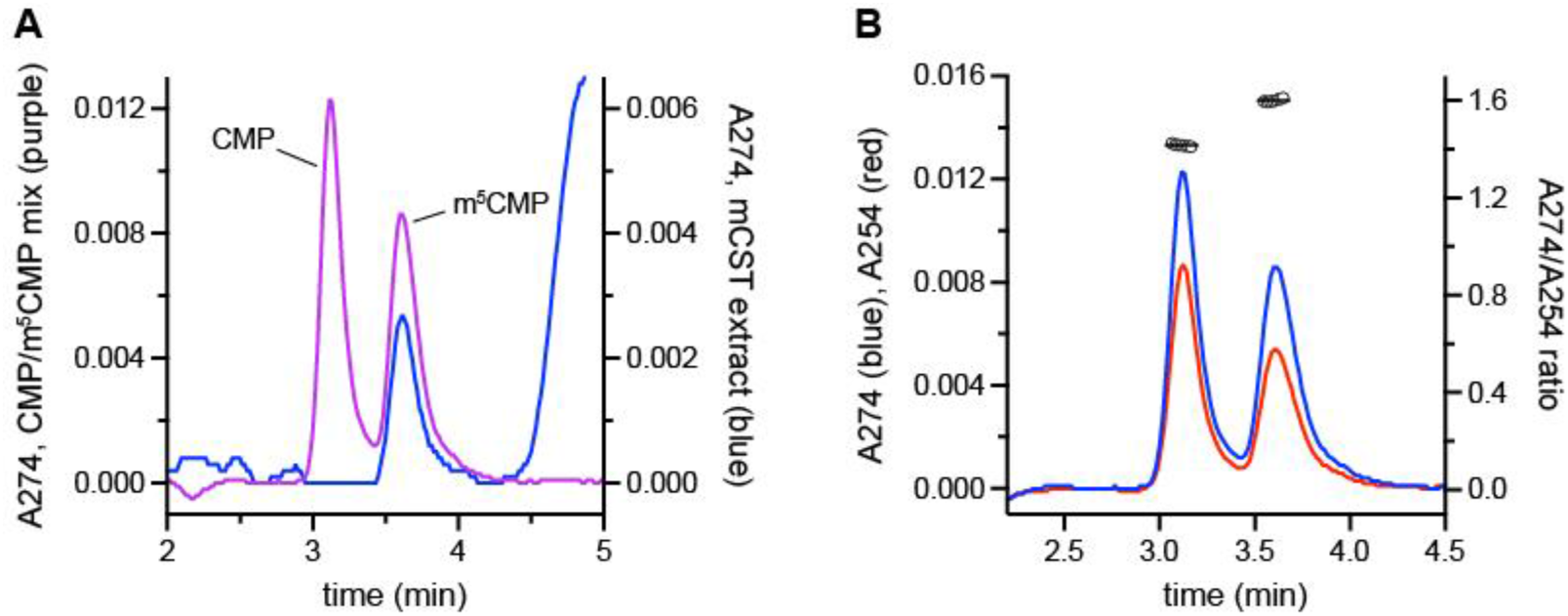
Comparison of m^5^CMP with the molecule that co-purifies with CST. **A)** The aqueous layer of a phenol-chloroform extraction of purified CST was run on a C18 HPLC column (blue trace). A separate sample containing 5 μM each of CMP and m^5^CMP was also run (purple trace). **B)** For the sample containing CMP and m^5^CMP, shown in panel A, the absorbance was monitored at both 274 nm (blue trace) and 254 nm (red trace) which is shown here. The ratio of the absorbance at these two wavelengths for a given time is plotted as the open circles on the right y-axis. The black horizontal lines represent the average A274/A254 ratio for the two peaks.

### m^5^CMP binds CST with a higher affinity than CMP and inhibits CMP-Sia uptake

We next wanted to characterize the functional properties of m^5^CMP and how it compares to CMP. The observation that m^5^CMP co-purifies with CST and remains bound even after overnight dialysis suggests that m^5^CMP has a slow off-rate, which would be most consistent with a sub-micromolar binding affinity. However, the assay that we have previously relied on to measure equilibrium binding constants is a scintillation proximity assay that requires 2 μM purified CST protein per assay point in order to achieve an adequate signal-to-noise ratio (Ahuja and Whorton 2019). Therefore, we thought it would be difficult to use this assay to measure m^5^CMP’s binding affinity towards CST since it would be very challenging to account for the significant ligand depletion that would occur.

So we instead developed an alternative assay that measures binding constants by evaluating the effect that a series of ligand concentrations has on the thermal stability of CST. In this assay, aliquots of 40 - 80 nM GFP-tagged CST are either kept at 4°C or heated to 41°C in the absence or presence of various concentrations of ligand. Some fraction of the CST protein will denature in response to the heating; however, the addition of a ligand will stabilize the protein and reduce the fraction of protein that denatures in a dose-dependent manner. The fraction of protein that remains folded can be determined by running the samples on a size exclusion column connected to a fluorescence detector and noting the peak height of the monodisperse species that elutes at ~5.4 min, as shown in Figure 4A.

**Figure 4.**
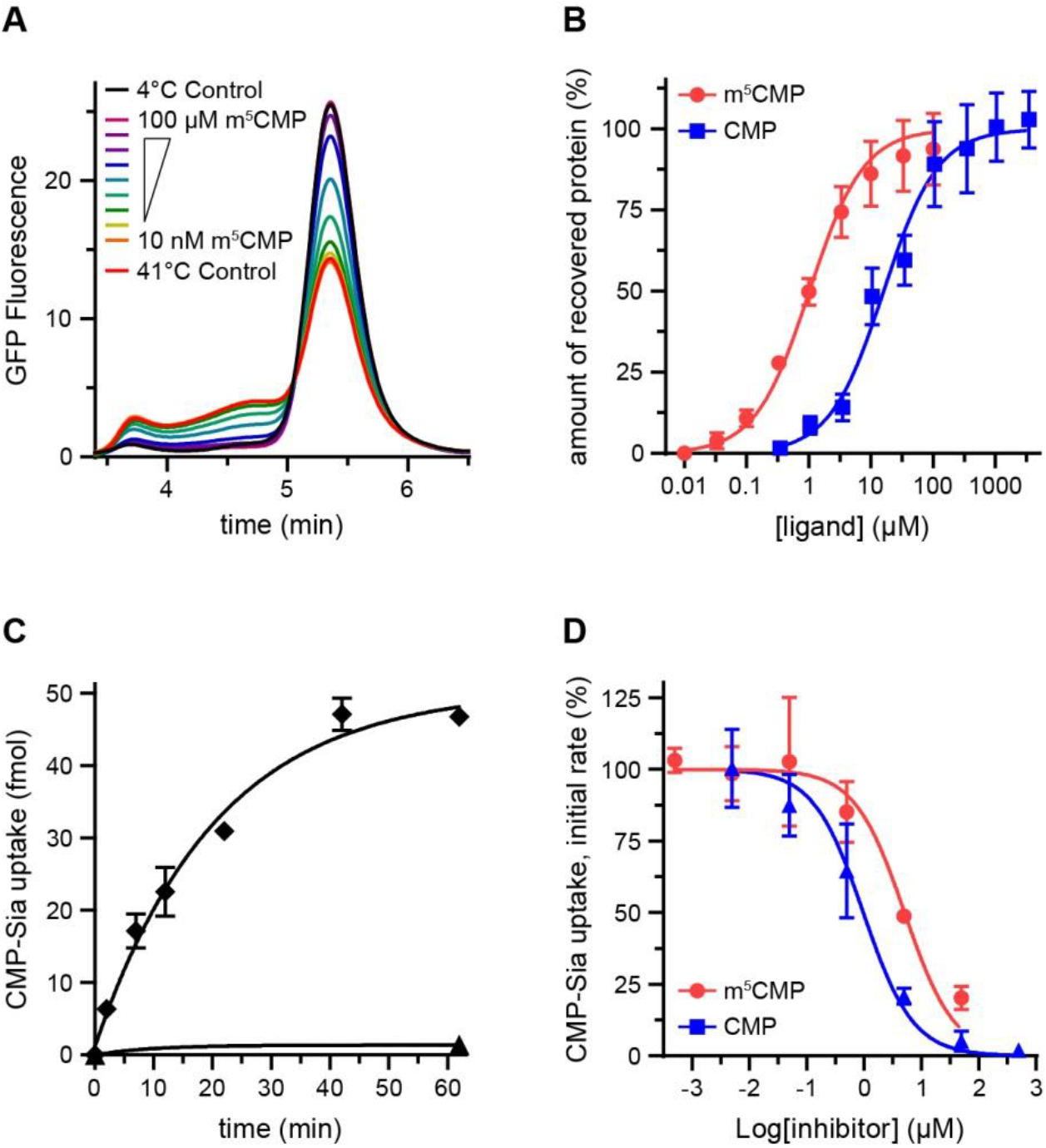
Binding constants and CMP-Sia transport inhibition for m^5^CMP and CMP. **A)** Aliquots of DDM-solubilized GFP-tagged CST protein were either kept at 4°C (black trace) or heated to 41°C, either alone (red trace) or in the presence of the indicated concentration of m^5^CMP for 15 min. The samples were clarified and then run on a gel filtration column with fluorescence detection. The sample eluting at ~5.4 min represents natively-folded protein. **B)** The peak heights from the traces in panel A were normalized to the highest (4°C with no ligand added) and lowest (41°C with no ligand added) peak height and are plotted against ligand concentration to determine binding constants. The plotted values represent the mean ± SEM, n=2. **C)** Intact Sf9 insect cells expressing CST (diamonds) or uninfected cells (triangles) were incubated with 30nM [^3^H]CMP-Sia for the indicated amount of time at room temperature. The plotted values represent the mean ± SEM, n=2. **D)** Intact Sf9 insect cells expressing CST (diamonds) were incubated with 30 nM [^3^H]CMP-Sia and the indicated concentration of inhibitor for 5 min at room temperature. The initial rate of uptake is plotted against inhibitor concentration in order to determine inhibition constants. The plotted values represent the mean ± SEM, n=4.

When we compare the peak heights of the CST sample heated to 41°C versus the sample kept at 4°C, we can see that approximately 44% of the protein denatures. However, including increasing amounts of m^5^CMP during the 41°C incubation leads to more and more protein being protected from denaturation, to the point where saturating amounts of m^5^CMP are able to prevent any denaturation – as indicated by the peak height for the 100 μM sample being identical to the sample that was kept at 4°C. By plotting peak heights against ligand concentration, we can determine a K_d_ of 1.0 ± 0.1 μM (Figure 4B). A similar experiment performed with a titration of CMP gives a K_d_ of 16.1 ± 2.4 μM.

We then wanted to compare m^5^CMP and CMP in their ability to inhibit CMP-Sia uptake. Again, our established vesicle-based *in vitro* transport assay requires micromolar amounts of protein per assay point (Ahuja and Whorton 2019, Cahill, Ahuja et al. 2020), so we developed a cell-based transport assay to reduce the amount of protein required. To do this, we first expressed CST in Sf9 insect cells using the baculovirus expression system. [^3^H]CMP-Sia was then incubated with intact cells at room temperature for the indicated time and the cells were then harvested by centrifugation before counting in a scintillation counter. As shown in Figure 4C, the rate of uptake is linear for at least 5 min. To measure inhibition constants, we added various concentrations of either m^5^CMP or CMP to the cells and incubated for 5 min. The amount of transport activity remaining as a function of ligand concentration is plotted in Figure 4D. Fitting the data with a simple dose-response model gives K_i_’s of 5.1 ± 1.2 μM and 1.0 ± 1.2 μM for m^5^CMP and CMP, respectively. The apparent discrepancy between these K_i_ values and the K_d_ binding constants will be discussed below.

### Structure of CST-m^5^CMP complex reveals the mechanism of high-affinity binding

In order to understand the molecular details of how m^5^CMP binds CST with a higher affinity than CMP, we determined the X-ray crystal structure of CST in complex with m^5^CMP. Crystals of CST were grown using the lipidic cubic phase method, in the presence of 400 μM m^5^CMP. Compared to crystals that we previously grew of CST in complex with CMP (Ahuja and Whorton 2019), crystals grown in the presence of m^5^CMP had the same morphology, belonged to the same space group, and had nearly identical unit cell properties (Table 1). However, one key difference is that the CST-m^5^CMP crystals diffracted X-rays to a much higher resolution of 1.8 Å (compared to 2.6 Å for the CST-CMP crystals). This let us build a highly-detailed model which contained 13 lipid molecules and 162 waters resulting in a R_work_ and R_free_ of 18.5% and 19.8%, respectively (Figure 5 – figure supplement 1 and Table 1).

**Table 1.**
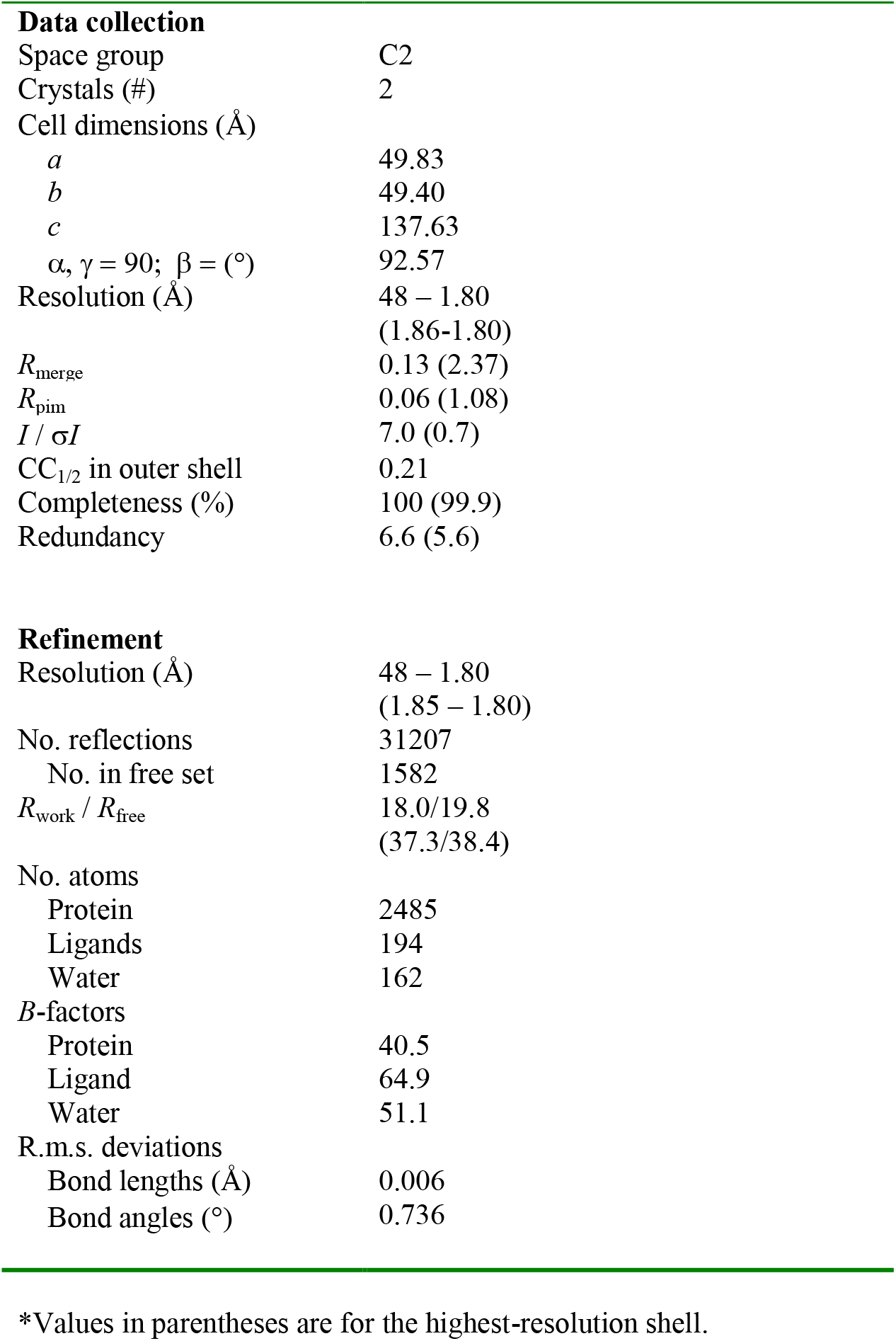
Data collection and refinement statistics

|  |  |
| --- | --- |
| **Data collection** |  |
| Space group | C2 |
| Crystals (#) | 2 |
| Cell dimensions (Å) |  |
| *a* | 49.83 |
| *b* | 49.40 |
| *c* | 137.63 |
| $\alpha, \gamma = 90; \beta = (^\circ)$ | 92.57 |
| Resolution (Å) | 48 – 1.80 |
|  | (1.86-1.80) |
| *R*<sub>merge</sub> | 0.13 (2.37) |
| *R*<sub>pim</sub> | 0.06 (1.08) |
| *I* / $\sigma I$ | 7.0 (0.7) |
| CC<sub>1/2</sub> in outer shell | 0.21 |
| Completeness (%) | 100 (99.9) |
| Redundancy | 6.6 (5.6) |
| **Refinement** |  |
| Resolution (Å) | 48 – 1.80 |
|  | (1.85 – 1.80) |
| No. reflections | 31207 |
| No. in free set | 1582 |
| *R*<sub>work</sub> / *R*<sub>free</sub> | 18.0/19.8 |
|  | (37.3/38.4) |
| No. atoms |  |
| Protein | 2485 |
| Ligands | 194 |
| Water | 162 |
| *B*-factors |  |
| Protein | 40.5 |
| Ligand | 64.9 |
| Water | 51.1 |
| R.m.s. deviations |  |
| Bond lengths (Å) | 0.006 |
| Bond angles (°) | 0.736 |

\*Values in parentheses are for the highest-resolution shell.

Overall, the CST-m^5^CMP structure is very similar to the CST-CMP structure, with a r.m.s.d. of 0.17 Å (Figure 5 – figure supplement 2). m^5^CMP binds in an essentially identical orientation as CMP does and there are no obviously-significant differences in the conformation of the residues that line the substrate-binding cavity (Figure 5A). The eponymous methyl group of m^5^CMP occupies a small and mostly-hydrophobic pocket (Figure 5B). This pocket is present but vacant in the CMP-bound structure (Figure 5C) which explains how CST can accommodate the addition of a methyl group at the C-5 position on the pyrimidine ring.

**Figure 5.**
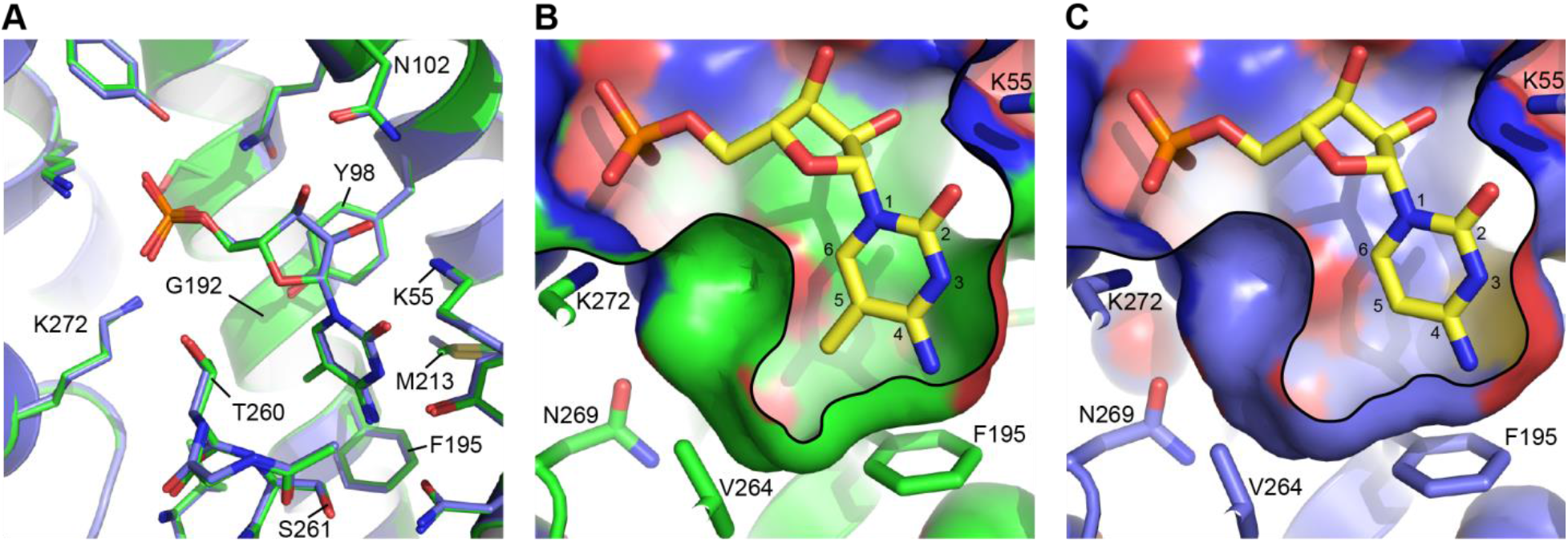
Comparison of the m^5^CMP and CMP binding sites. **A)** A close-up of the substrate binding pocket is shown for both the CST-m^5^CMP (green) and CST-CMP (blue) structures. Only substrate-interacting side chains are shown and portions of TM8 are hidden for clarity. **B and C)** A slice through the surface representation of the substrate-binding cavity is shown for the CST-m^5^CMP (B) and CST-CMP (C) structures. The atoms for the pyrimidine ring are numbered. A black line is added as a visual aide to indicate where the cavity volume was sliced. The view is similar to that shown in panel A but is adjusted slightly to better show the part of the substrate-binding cavity that interacts with the C-5 methyl.

The burial of m^5^CMP’s C-5 methyl in this hydrophobic pocket is likely the primary contributor to the 16-fold increase in m^5^CMP’s binding affinity compared to CMP, on account of the hydrophobic effect. It has been estimated that burying hydrophobic surfaces contributes approximately 0.03 kcal/mol/Å^2^ to the free energy of ligand binding (Hopkins, Chothia 1974). Therefore, burying a methyl group, which has a surface area of 46 Å^2^, would contribute a total of about 1.4 kcal/mol which is equivalent to a ~10-fold increase in affinity. This general effect has been termed the magic methyl effect (Schonherr and Cernak 2013) and it is not uncommon to see such large increases in binding affinity under the right circumstances – *e.g*. where an added methyl group is buried in a hydrophobic pocket (Leung, Leung et al. 2012).

The residues that line the methyl-interacting hydrophobic pocket – primarily Phe195, Thr260 (C^γ2^ atom), and Val264, but also Tyr98, Gly192, Met213, and Ser261 (only C^α^ and C^β^ atoms) (Figure 5) are highly conserved among orthologous CST proteins (SLC35A1 gene products) from other organisms (Figure 5 – figure supplement 3), suggesting that the ability for CST to bind m^5^CMP is a conserved property across species. Many of these substrate-binding cavity residues are also conserved in the homologous UGT and NGT proteins (SLC35A2 and SLC35A3 gene products, respectively; Figure 5 – figure supplement 3), which transport UMP and several UDP-coupled sugars (Ishida and Kawakita 2004, Hadley, Maggioni et al. 2014). As shown in Figure 5 – figure supplement 4, docking UMP into structural homology models of UGT and NGT suggest that these proteins also contain a vacant and mostly hydrophobic pocket adjacent to the C-5 position of the uracil group of UMP. This raises the question of if UGT or NGT can also interact with m^5^CMP or other UMP-like nucleotide analogs, such as pseudouridine or methyluridine monophosphates, that are also common epigenetic modifiers of RNA.

## Discussion

Here we describe the discovery that m^5^CMP binds CST and can inhibit CMP-Sia transport. Using a thermal shift assay, we determined that m^5^CMP has a binding constant of 1 μM, which implies a relatively fast off-rate and short half-life on the order of a second or less. However, in order to co-purify a molecule over the course of a 2-day protein purification protocol, a relatively slow off-rate with a half-live on the order of at least several hours would be required. This would approximately equate to a low-nanomolar dissociation constant. The discrepancy between these two dissociation constants can be reconciled by the fact that the binding experiment was performed at the T_m_ of CST (41°C) whereas the protein purification was primarily performed at 4°C. This suggests that there is a steep relationship between temperature and m^5^CMP binding affinity. It is not straightforward to extrapolate a K_d_ at intermediate temperatures; however, it follows that the K_d_ for m^5^CMP at lower temperatures, such as a physiological temperature of 37°C or room temperature where transport assays are performed, will be lower than what was observed at 41°C – perhaps on the order of several hundred nanomolar.

Another seemingly paradoxical finding is that although m^5^CMP has a lower binding K_d_ than CMP, the apparent K_i_ for m^5^CMP’s inhibition of CMP-Sia transport is about 5-fold higher than that of CMP (5.1 μM versus 1.0 μM). The transport assay was performed at room temperature, so it is understandable that the K_i_ for CMP is lower than the K_d_ of 16 μM that we measured at 41°C using the thermal shift binding assay. While K_i_’s of transport inhibition and K_d_’s of binding are not always necessarily identical, they are often quite similar (Van Winkle 1999). In fact, this does actually seem to be the case for CMP inhibition of CST, considering that we previously determined that the K_d_ for CMP binding to CST was on the order of 1-6 μM at room temperature using a scintillation proximity binding assay (Ahuja and Whorton 2019). Therefore, we expected that the relationship between the K_i_’s for m^5^CMP and CMP would have mirrored what we observed for their K_d_’s. In other words, we would have expected to see that the K_i_ for m^5^CMP inhibition of CMP-Sia transport to be significantly lower than CMP’s K_i_.

The reason that we did not observe this is not entirely clear. It may be the case that m^5^CMP’s K_i_ is indeed higher than CMP’s K_i_ despite having a lower equilibrium dissociation constant. Given that CSTs transport mechanism likely involves several conformational states (Ahuja and Whorton 2019), it is possible that in steady-state conditions m^5^CMP’s extra methyl adversely affects its interaction with certain conformational states or affects the rates of transitions between states. There could also be a number of technical reasons that may underlie the discrepancy between m^5^CMP’s binding K_d_ and transport K_i_. The transport assay relies on intact cells, so it is possible that there are cellular processes that either preferentially degrade or uptake m^5^CMP over CMP, thereby affecting its effective concentration. In addition, m^5^CMP’s additional methyl group imparts significant hydrophobicity making it at least 100-fold less soluble in water. Therefore, given that we use a large number of cells per assay point (1 x 10^6^), this presents a significant amount of lipid bilayer where the m^5^CMP may partition which could lower the effective concentration of soluble m^5^CMP. Finally, as mentioned above, our goal for this assay was to reduce the total concentration of transporter in the assay; however, it may be the case that the concentration of CST per assay point is still too high such that ligand depletion affects our ability to accurately measure sub-μM K_i_’s. To address these technical concerns, future work will need to characterize the actual concentration of soluble extracellular m^5^CMP as well as define the concentration of CST per assay point.

There are two requirements for m^5^CMP to act as a physiological inhibitor of CMP-Sia transport: 1) physiological concentrations of m^5^CMP should align with m^5^CMP’s inhibition binding constants, and 2) physiological ratios of m^5^CMP and CMP concentrations should be similar to the observed ratios in binding constants. Until recently, it has not been possible to detect the levels of m^5^CMP inside cells; however, a recent development of a sensitive detection method has provided the first insight into cellular m^5^CMP concentrations (Zeng, Qi et al. 2017). In this work, Zeng et al. measured the concentration of a panel of nucleotides, including CMP, m^5^CMP, and deoxy-m^5^CMP (m^5^dCMP) for the HEK293T and HeLa cultured cell lines as well as for a number of samples of human renal tissues. They found that CMP concentrations in these samples ranged from 6.7 - 72.6 pmol/mg protein whereas m^5^CMP concentrations ranged from 0.004 - 0.02 pmol/mg protein, which were typically several fold higher than m^5^dCMP levels. This equates to ratios of [CMP]:[m^5^CMP] ranging from 690 to 7260. Molar concentrations of CMP in renal tissues have not been reported, but CMP molarities in several other mammalian tissues have been reported, with CMP concentrations ranging from 4.8 - 96 μM with an average of 39±2 μM. This would translate to an approximate molar concentration for m^5^CMP of between 5 - 60 nM. This low concentration, as well as the presence of a vast excess of CMP, most likely means that m^5^CMP would not be concentrated enough to affect CST transport activity.

However, Zeng et al. went further and also measured the concentrations of the same panel of nucleotides from human urine samples. Although the concentrations cannot be compared to cellular concentrations because they were determined in terms of pmol/mg creatinine, it is still valid to compare the ratio of [CMP]:[m^5^CMP], which was about an order of magnitude lower than what was seen in the renal tissue samples, ranging from 56 - 486. This indicates that not only is m^5^CMP widely distributed in the body, but there are likely cell types where m^5^CMP is much more abundant, compared to CMP levels, than what was seen in renal cells. In addition, the m^5^CMP levels detected in urine is a conglomeration from all tissue types; therefore, since there are tissues types with high CMP:m^5^CMP ratios (e.g. renal), it follows that there must be some tissue types with CMP:m^5^CMP ratios even lower than the averages that were observed in the urine samples – perhaps even approaching the levels mirroring the 16-fold affinity difference that we measured between CMP and m^5^CMP. Using the same estimate for an average cellular CMP concentration of 39 μM, this would be equivalent to m^5^CMP concentrations ranging from 80 - 700 nM. Therefore, it is conceivable that there are cell types, perhaps ones that have high RNA turnover and/or are predisposed to high levels of RNA cytosine methylation, where the physiological levels of m^5^CMP would be adequate to regulate the transport activity of CST.

This analysis has focused on m^5^CMP since this was the molecule that we identified to co-purify with CST; however, as mentioned above, m^5^dCMP is also present in cells. In the Zeng et al. study, m^5^dCMP concentrations were observed to typically be approximately several fold lower than m^5^CMP concentrations, ostensibly because m^5^dCMP originates from DNA turnover which in most cell types happens less frequently than RNA turnover. Deoxy-cytidine differs from cytidine by the removal of the 2’ hydroxyl from the ribose group. Both this 2’ hydroxyl as well as the 3’ hydroxyl form hydrogen bonds with Asn102 of CST, as well as with surrounding structured waters (Figure 5A, figure 5 – supplement 1D, and Ahuja and Whorton ^20^). However, it does not appear that the loss of the 2’ hydroxyl significantly affects binding affinity since it has been previously shown that CMP and dCMP have essentially identical K_i_’s for inhibition of CMP-Sia transport (Chiaramonte, Koviach et al. 2001). Therefore it seems that cellular pools of m^5^dCMP may also be able to contribute to inhibition of CMP-Sia transport.

In conclusion, we have shown that m^5^CMP co-purifies with CST and most likely represents a novel physiological regulator of CST transport activity. This work has focused on characterizing the interaction between m^5^CMP and the mouse ortholog of CST. However, considering the very high sequence identity between the mouse and human CST sequences (Figure 5 – figure supplement 3), especially in regards to residues that line the substrate-binding pocket, we expect that human CST will have nearly identical properties as mouse CST. m^5^CMP binds CST with a 16-fold higher equilibrium binding affinity than CMP, but m^5^CMP’s K_i_ for inhibition of CMP-Sia transport is paradoxically approximately 5-fold higher than that of CMP. However, we discuss how there may be several technical reasons for this discrepancy. If m^5^CMP’s K_i_ is also roughly 16-fold lower than CMP’s K_i_, mirroring what was observed for the binding K_d_’s, then there are likely some cell types where the cellular m^5^CMP concentration is high enough to approach m^5^CMP’s K_i_. In these cases, fluctuations in m^5^CMP and m^5^dCMP concentrations that are connected to rates of RNA/DNA cytosine methylation and decay would be able to impact the uptake of CMP-Sia into the Golgi lumen and thereby affect glycosylation patterns. Ultimately, experiments that can monitor glycosylation profiles in response to manipulation of rates of cellular RNA/DNA methylation and/or decay will be crucial for establishing a definitive link between RNA/DNA epigenetics and regulation of glycosylation through alteration of NST activity.

## Materials and Methods

### Protein expression and purification

Expression and purification of CST was performed as previously described (Ahuja and Whorton 2019, Cahill, Ahuja et al. 2020). Briefly, the full-length mouse CST construct (with a C-terminal PreScission protease site, followed by green fluorescent protein (GFP), and then a His10 tag) was expressed in *P. pastoris* cells. The cells were lysed by cryogenic milling and then solubilized using the detergent n-dodecyl-β-D-maltopyranoside (DDM) (Anatrace, solgrade). CST was purified from the clarified lysate using Talon resin (Clontech) followed by protease cleavage of the GFP-His10 tag and finished with size exclusion chromatography on a Superdex 200 column equilibrated in Buffer A (25 mM HEPES pH 7.5, 150 mM NaCl, 0.1% (w/v) DDM (anagrade), 5 mM DTT, and 1 mM EDTA).

### HPLC analysis

HPLC analysis was performed as previously described (Ahuja and Whorton 2019). Briefly, samples were run on a XSelect CSH C18 column (3.5 μm, 2.1 × 150 mm; Waters) using a mobile phase of Buffer B (0.1 M K•PO_4_ pH 6.5, 8 mM tetrabutylammonium hydrogensulfate (TBHS)). For samples containing known compounds, stocks of the compound were diluted into Buffer B. For samples containing protein extracts, a volume of typically 50 μM purified CST in Buffer A was added to an equal volume of a phenol-chloroform-isoamyl alcohol mix (25:24:1) in a 1.5 ml microcentrifuge tube. This mixture was vigorously vortexed for 1 min and then spun at 21,000 x g for 5 min in a microcentrifuge. The top aqueous layer was collected and added to an equal volume of chloroform. This was again vortexed for 1 min and then spun at 21,000 x g for 5 min in a microcentrifuge. The top aqueous layer was collected for HPLC analysis. For AnP-treatment of CST prior to phenol-chloroform extraction, 3 μl AnP stock (5,000 units/ml, New England Biolabs) was added to 50 μM purified CST in Buffer A along with 0.5 mM ZnCl_2_ and 1mM MgCl_2_. For protein samples that were dialyzed prior to phenol-chloroform extraction, 100 μl aliquots of 50 μM purified CST were placed in a dialysis cassette with a 10K MWCO and dialyzed against 100 ml at 4°C with at least three separate buffer exchanges over the course of 24 hr.

### Mass Spectrometry

The HPLC-ESI-MS measurements were carried out by the Northwest Metabolomics Research Center at the University of Washington using an Agilent 6545 Q-TOF mass spectrometer coupled with an Agilent 1290 Infinity LC pump, and an Agilent 6520 Q-TOF mass spectrometer coupled with Agilent 1260 Infinity LC system (Agilent Technologies). Samples consisted of phenol-chloroform extracts (prepared as described above) of either Buffer A alone or 50 μM purified CST in Buffer A. The HPLC separation was performed using a Waters XBridge BEH Amide column (15 cm x 2.1 mm, 2.5 μm). The mobile phase consisted of (A) H_2_O:acetonitrile (95:5, v/v), 5 mM ammonium acetate, and 0.1% acetic acid, and (B) H_2_O:acetonitrile (5:95, v/v), 5 mM ammonium acetate, and 0.1% acetic acid. Gradient operation was initiated at 94% of solvent B, and it decreased to 78% at t = 6.5 min, and to 39% at t = 12.0 min. Composition was maintained at 39% of solvent B until t = 18.5 min, followed by an increase to 94% at t= 19.0 min, and maintained at this condition until t = 35.0 min (total experimental time for each injection). The flow rate was 0.3 ml/min, the injection volume was 5 μl, followed by H_2_O:acetonitrile (5:95, v/v) needle wash for 10 s. The column was maintained at 35°C. The ESI conditions were as follows: electrospray ion source ESI Agilent Jet Stream Technology in positive ionization mode; voltage 3.8 kV; desolvation temperature 325°C; cone flow 20 l/h; desolvation gas flow 600 l/h; nebulizer pressure 45 psi, N_2_ was used as drying gas; MS scan rate of 5 spectra/s across the range *m/z* 60-1000, threshold count: 200; MS/MS acquisition rate 3 spectra/s, targeting precursor ion *m/z* 338.0701. Isolation width of 1.3 *m/z*, collision energies of 10 and 15 eV, *m/z* range of 50-500. Data were acquired using MassHunter Data Acquisition Workstation v. B.06.01.6157 software (Agilent Technologies, Palo Alto, CA). All solvents were LC-MS grade (Fisher Scientific). DI water (18.2 MΩ•cm at 25°C) was provided in-house by a Synergy Ultrapure Water System (EMD Millipore).

### Thermal shift binding assay

Full-length, GFP-tagged mouse CST was expressed in *P. pastoris* as described above. Milled cells were suspended in lysis buffer at a ratio of 125 mg cells to 1 ml buffer, then gently rotated for 2 hours at 4°C (lysis buffer: 50 mM HEPES pH 7.5, 150 mM NaCl, 1% w/v DDM (Anatrace, solgrade), 1 mM DTT, 1 mM EDTA, 0.01 mg/ml deoxyribonuclease I, 0.7 μg/ml pepstatin, 1 μg/ml leupeptin, 1 μg/ml aprotinin, 1 mM benzamidine, 0.5 mM phenylmethylsulfonyl fluoride). This DDM-solubilized lysate was then clarified by centrifugation at 21,000 x g for 20 minutes at 4°C. It was then diluted 32-fold in Buffer C (50 mM HEPES pH 7.5, 150 mM NaCl, 0.1% w/v solgrade DDM, 1 mM EDTA, 1 mM DTT). This dilution factor was chosen to give a final assay concentration of 40-80 nM GFP-tagged CST and was based on comparing the peak heights of the 4°C control peak with samples of previously-run purified GFP-tagged CST of known concentration (data not shown). 90 μl aliquots of the diluted lysate were then placed into 250 μl PCR tubes (Fisher Scientific). 10 μl of 10X stocks of either CMP or m^5^CMP (made up in 25 mM HEPES pH 7.5 and 150 mM NaCl) were then added to the diluted lysate samples. The samples were gently mixed and incubated on ice for 30 minutes. Following this, they were then heated to 41°C for 15 minutes using a thermocycler (control samples were kept on ice), transferred to 1.5 ml microcentrifuge tubes, and spun down at 87,000 g for 20 minutes at 4°C to pellet precipitated protein and cellular debris.

The supernatants were analyzed by size exclusion chromatography coupled to fluorescence detection (FSEC). Briefly, supernatants were transferred into the wells of a 96-well sample block. A Waters Acquity UPLC with a fluorescence detector was used to apply 50 μL of each sample to a Superdex 200 Increase GL 5/150 column equilibrated in Buffer C. Each sample was analyzed for GFP fluorescence during 10 minutes of chromatography at a flow rate of 0.3 ml/min. Peak heights from individual FSEC runs were normalized to the difference between the highest (4°C with no substrate added) and lowest (41°C with no substrate added) peak height, and fitted to a single-site binding model using Prism (GraphPad).

### Cell Transport Assay

Sf9 insect cells (Expression Systems) were grown in suspension in ESF 921 media (Expression Systems) to a density of 1×10^6^ cells per ml and then infected with baculovirus encoding the same full-length mouse CST construct as described above at a ratio of 40 μL virus per ml of media (this ratio was empirically determined from titer trials in order to optimize protein expression). Cells were allowed to express the protein for 48 hours, counted, centrifuged at 1000 x g for 5 minutes at room temperature, and resuspended in fresh media to a density of 2×10^6^ cells/ml. Aliquots of 0.5 ml (1×10^6^ cells) were made in 1.5 ml microcentrifuge tubes. Uninfected cells were similarly counted, spun down, resuspended, and aliquoted for use as control samples.

For time-course CMP-Sia uptake experiments, [^3^H]CMP-Sia (20 Ci/mmol; American Radiolabeled Chemicals) was added to each sample for a final concentration of 30 nM. The tubes were gently rotated at room temperature for their designated incubation times, then centrifuged at 1000 x g for 3 minutes. Each sample underwent three cycles of washing in 500 μl ice-cold PBS followed by another round of centrifugation. Finally, the pellet was resuspended in 200 μl of PBS, transferred to a 7 ml scintillation vial with 5 ml scintillation fluid (Ultima Gold, PerkinElmer), inverted several times to mix, and counted for 3 minutes in a scintillation counter. The specific counts for each trial were determined by subtracting CPMs of the uninfected control cells from those of the corresponding infected cells.

For inhibitor titration experiments, 100x stock concentrations of inhibitor (either CMP or m^5^CMP) were made in PBS. Inhibitor stock (5 μl) or PBS (5 μl; for uninhibited control samples) was added to each tube of freshly resuspended cells, which were then gently inverted several times to mix and allowed to incubate for 5 minutes at room temperature. [^3^H]CMP-Sia was added to a final concentration of 30 nM. The samples were rotated at room temperature for 5 minutes, and then centrifuged, washed, and counted as described above.

### Crystallography and structure determination

For crystallization of CST in complex with m^5^CMP, the CSTΔC construct, which lacks the last 15 residues (322-336), was expressed and purified as previously described (Ahuja and Whorton 2019). However, before the final size exclusion chromatography step, AnP was added to the Talon-purified protein and dialyzed overnight to remove any bound m^5^CMP. After the final size exclusion chromatography and concentration, m^5^CMP was added to the protein at 400 μM and crystallized as previously described (Ahuja and Whorton 2019). Briefly, the protein was mixed 2:3 with monoolein (Nu-Chek Prep) and then 70 nl of this material was deposited on a glass slide (Molecular Dimensions). 600 nl of the crystallization solution (26.7–30% PEG 300, 0.1 M MES pH 6.5, and 0.1 M NaCl) was then added. The drop was sealed with a glass slide on the top and incubated at 20°C. The crystals were then harvested directly from the drop and flash-frozen in liquid N_2_.

Diffraction data were collected at the APS 23ID-D beamline using an X-ray wavelength of 1.03319 Å. The data were processed with XDS (Kabsch 2010) and further analyzed using Pointless (Evans 2011) and Aimless (Evans and Murshudov 2013). CCP4i (Winn, Ballard et al. 2011) was used for project and job organization. The structure was solved by molecular replacement using Phaser (McCoy, Grosse-Kunstleve et al. 2007), using the CST-CMP crystal structure model (PDB id: 6OH2 (Ahuja and Whorton 2019)) as a search model. Iterative cycles of model building in Coot (Emsley, Lohkamp et al. 2010) and refinement using Refmac (Murshudov, Skubak et al. 2011) and Phenix (Adams, Afonine et al. 2010) were used to add in m^5^CMP, build the previously unmodeled loop connecting TMs 5&6, and build the last 10 residues of the construct (321-330) which is mostly comprised of PreScission protease recognition site. The “Find Waters” feature in Coot was used to automatically identify structured waters which were then manually inspected. The final model consists of residues 9-330 (minus 318-320), 162 waters, 13 monooleins, and 2 PEGs. The model was validated using MolProbity (Chen, Arendall et al. 2010) as implemented in Phenix. The model had 98.7% of its residues in the preferred region of a Ramachandran plot and no outliers. Figures were prepared using PyMOL (Schrodinger 2015).

### Homology modeling

Structural homology models of UGT and NGT bound to UMP were generated as previously described (Ahuja and Whorton 2019). Briefly, the models were generated using the SWISS-MODEL web server (Waterhouse et al., 2018), with the mCST-CMP-Sia structure used as a template. UMP was then placed into these models in the same pose that CMP is found in the mCST-CMP-Sia structure.

### Data availability

Atomic coordinates and structure factors for the CST-m^5^CMP structure have been deposited in the Protein Data Bank (PDB) with an entry ID of 6XBO.

## Acknowledgements

We thank the GM/CA (APS) beamline staff, M. Becker, N. Sanishvili, and N. Venugopalan, for assistance with remote data collection. We thank D. Raftery, N. Gowda, and F. Carnevale of the Northwest Metabalomics Research Center for help in analyzing and interpreting the mass spectrometry data. We thank E. Gouaux for use of crystallization robotics and scintillation counters. This work was supported by NIH grant R01GM130909.

---

**Figure 2 – figure supplement 1.**
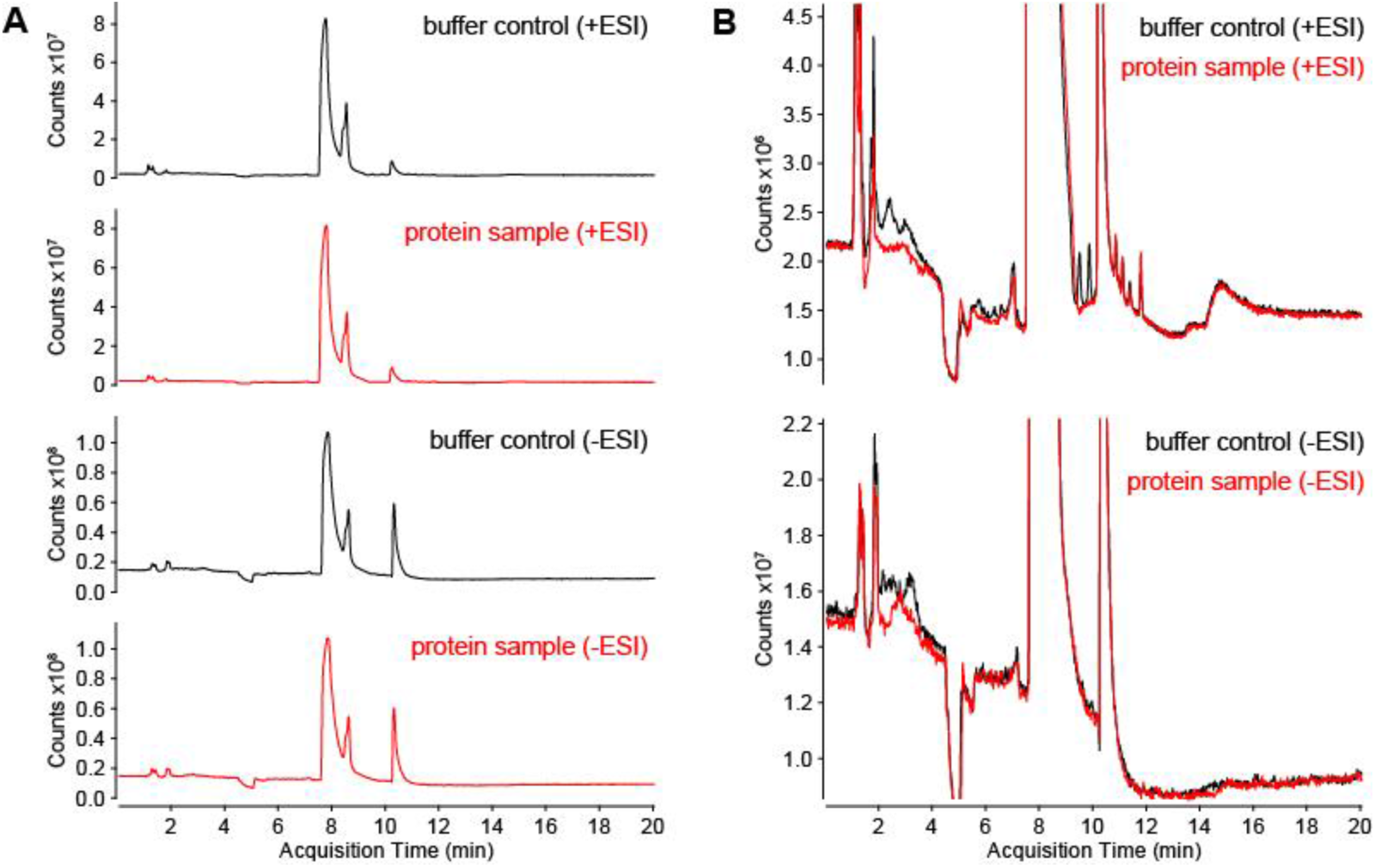
Comparison of total ion current chromatograms for buffer-only versus CST protein-containing samples. Phenol-chloroform extractions were performed on either a buffer-only control (black traces) or a sample containing purified CST protein (red traces). The aqueous layers were run on an LC-MS system in either positive or negative ESI mode with the resulting total ion current chromatograms shown in panel A. The chromatograms for each ESI mode are overlaid for better comparison in panel B.

**Figure 5 – figure supplement 1.**
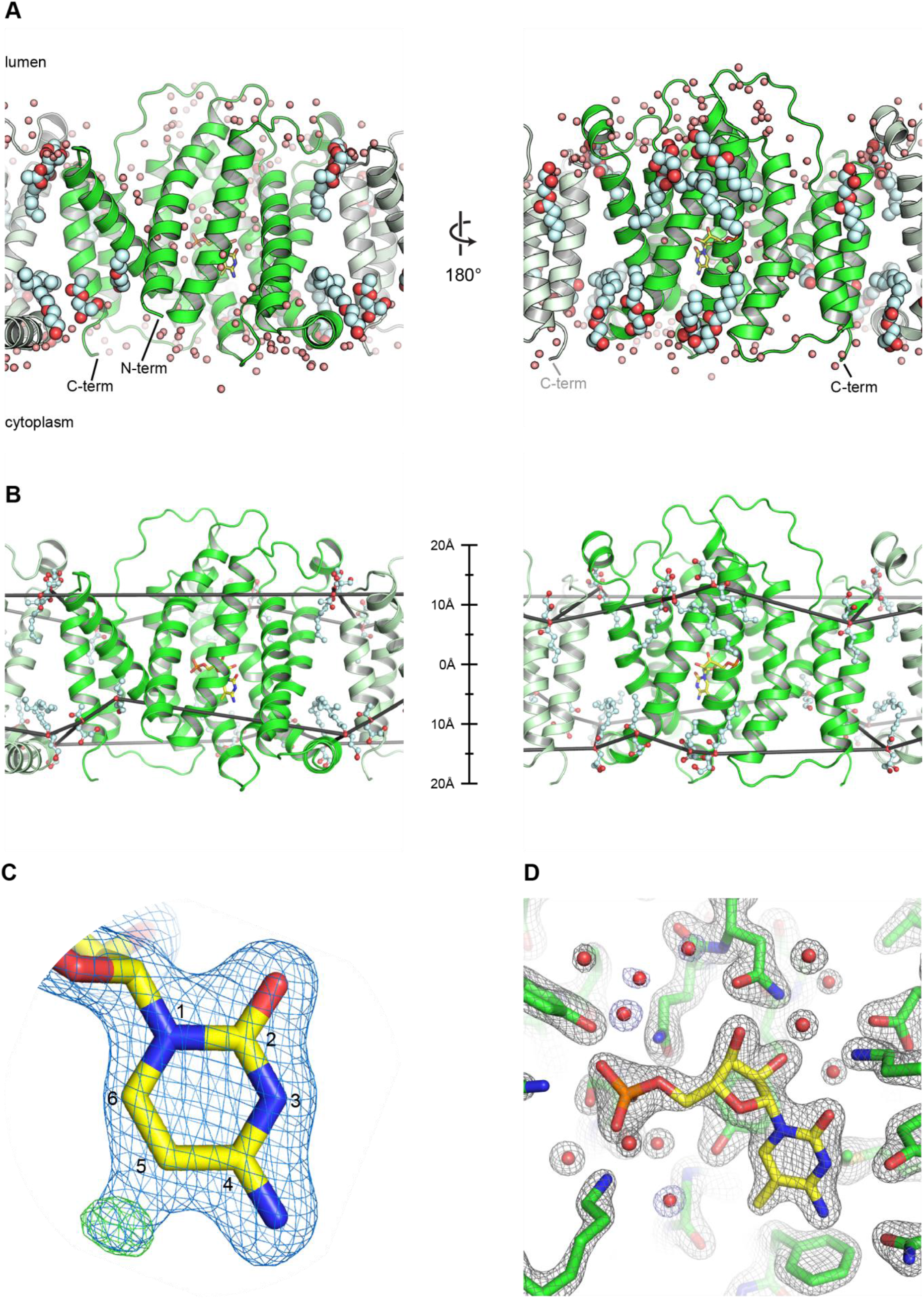
Details of water, lipid, and substrate modeling for the CST-m^5^CMP structure. **A)** Two views (front and back) of the CST-m^5^CMP structure are shown, with the lumenal and cytoplasmic sides of the protein indicated. Water molecules are shown as small pink spheres and monoolein lipid molecules are shown as larger spheres with red indicating oxygen and light cyan indicating carbon. The CST molecule in the asymmetric unit is shown as a green cartoon. Symmetry-related CST molecules to the left and right are shown (as light green cartoons) since some of the lipids and waters mediate crystal contacts between CST molecules. **B)** The same views of the CST-m^5^CMP structure that are shown in panel A are show here as well, except only the monoolein lipid molecules are shown in ball-and-stick representation. The oxygen atoms of the acyl-ester linkages of adjacent monooleins are connected by a black line to give a first order approximation of the shape and thickness of a lipid bilayer that would interact with CST. A scale bar is shown to indicate the extent of the protein-lipid interface on either side of the protein. **C)** 2F_o_-F_c_ (blue mesh, 1.5σ) and F_o_-F_c_ (green mesh, 3.5σ) electron density maps are shown. These maps were obtained by using molecular replacement to solve the structure of the CST-m^5^CMP crystal, using the CST-CMP structure as a search model. The cytosine group of CMP from the search model is shown as yellow sticks. The atoms of the pyrimidine ring are numbered and density for a methyl group at the C-5 position is clearly observed in both maps. **D)** A 2F_o_-F_c_ map of the final refined structure of the CST-m^5^CMP structure is shown. The gray mesh is contoured at 1.8σ and the blue mesh is contoured at 1σ to show the weaker density for some of the waters.

**Figure 5 – figure supplement 2.**
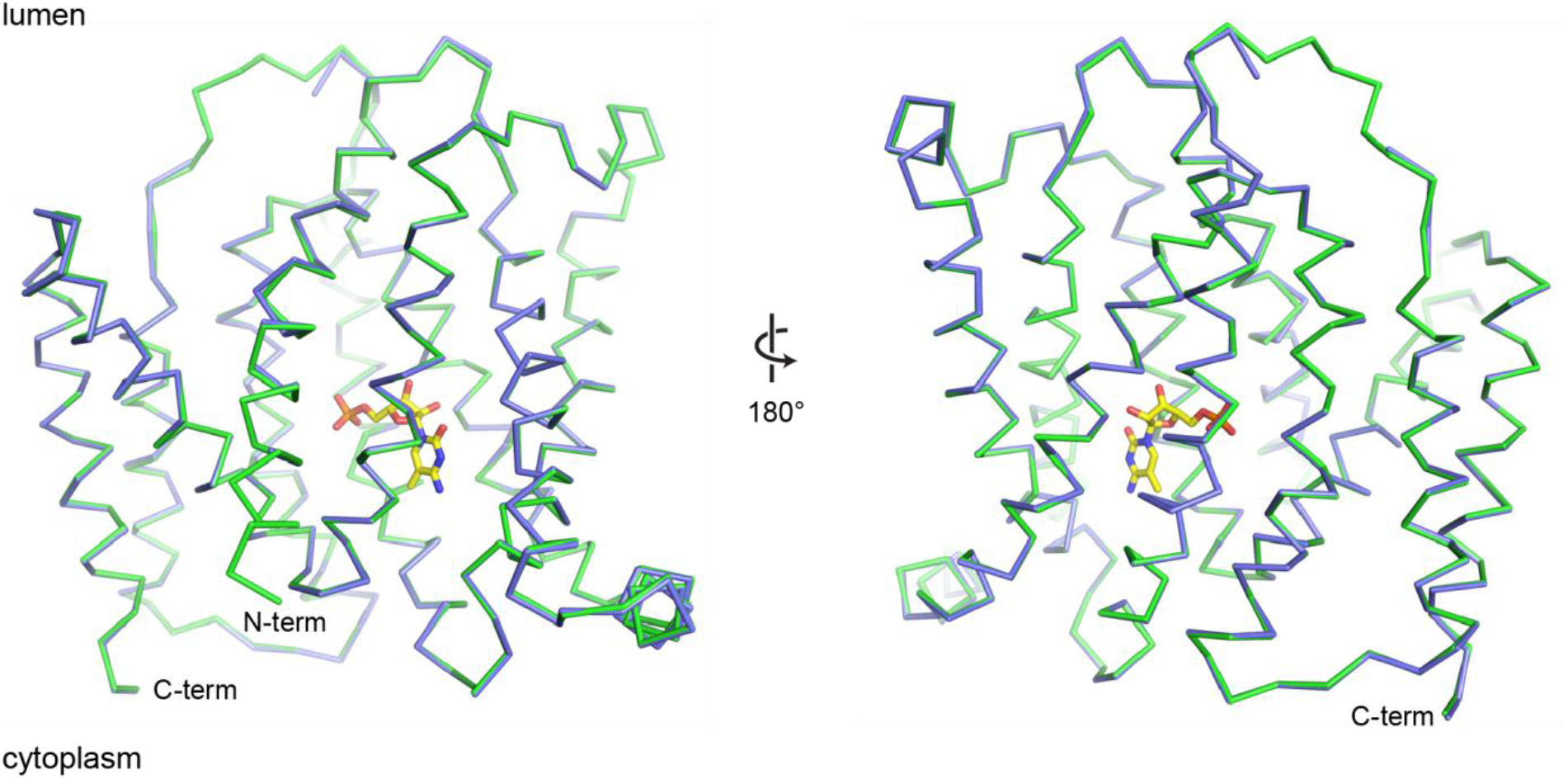
Comparisons of the CST-CMP and CST-m^5^CMP structures. The CST-CMP (blue) and CST-m^5^CMP (green) structures are shown as Cα traces and are superimposed to show the high structural identity. Front and back views are shown with the lumenal and cytoplasmic sides of the protein indicated. m^5^CMP is shown as yellow sticks.

**Figure 5 – figure supplement 3.**
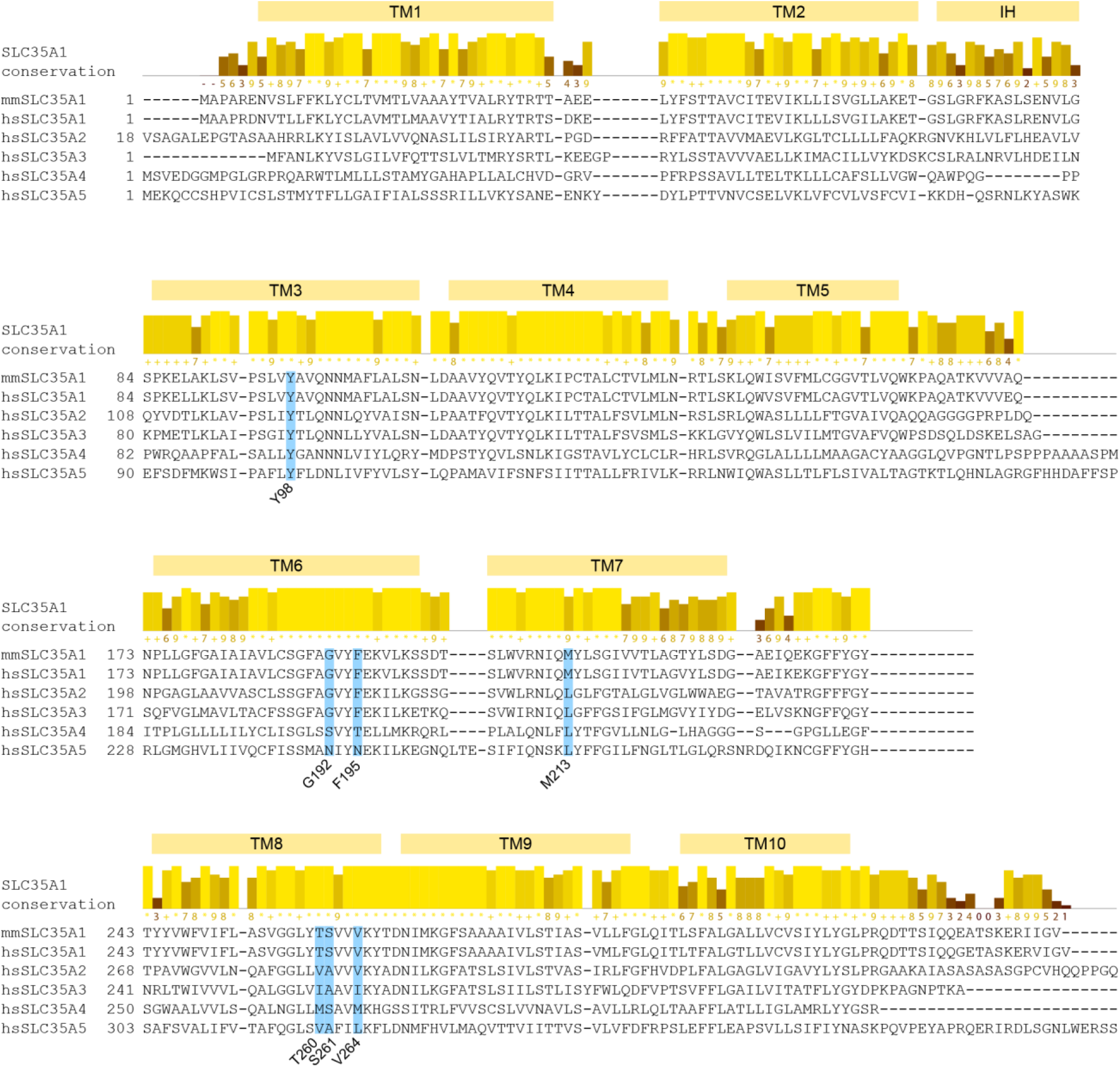
SLC35A sequence conservation and alignment. A sequence alignment between mouse CST (mmSLC35A1), human CST (hsSLC35A1), and other human SLC35A family members is shown. The bar graph above the alignment shows the sequence conservation among 126 SLC35A1 orthologs. The row of numbers and symbols under the bar graph indicates the degree of conservation, with a higher number indicating greater conservation, a “+” symbol indicating near-complete identity, and an “*” symbol indicating complete identity. Residues relevant to m^5^CMP interactions that are discussed in the text are highlighted and labeled.

**Figure 5 – figure supplement 4.**
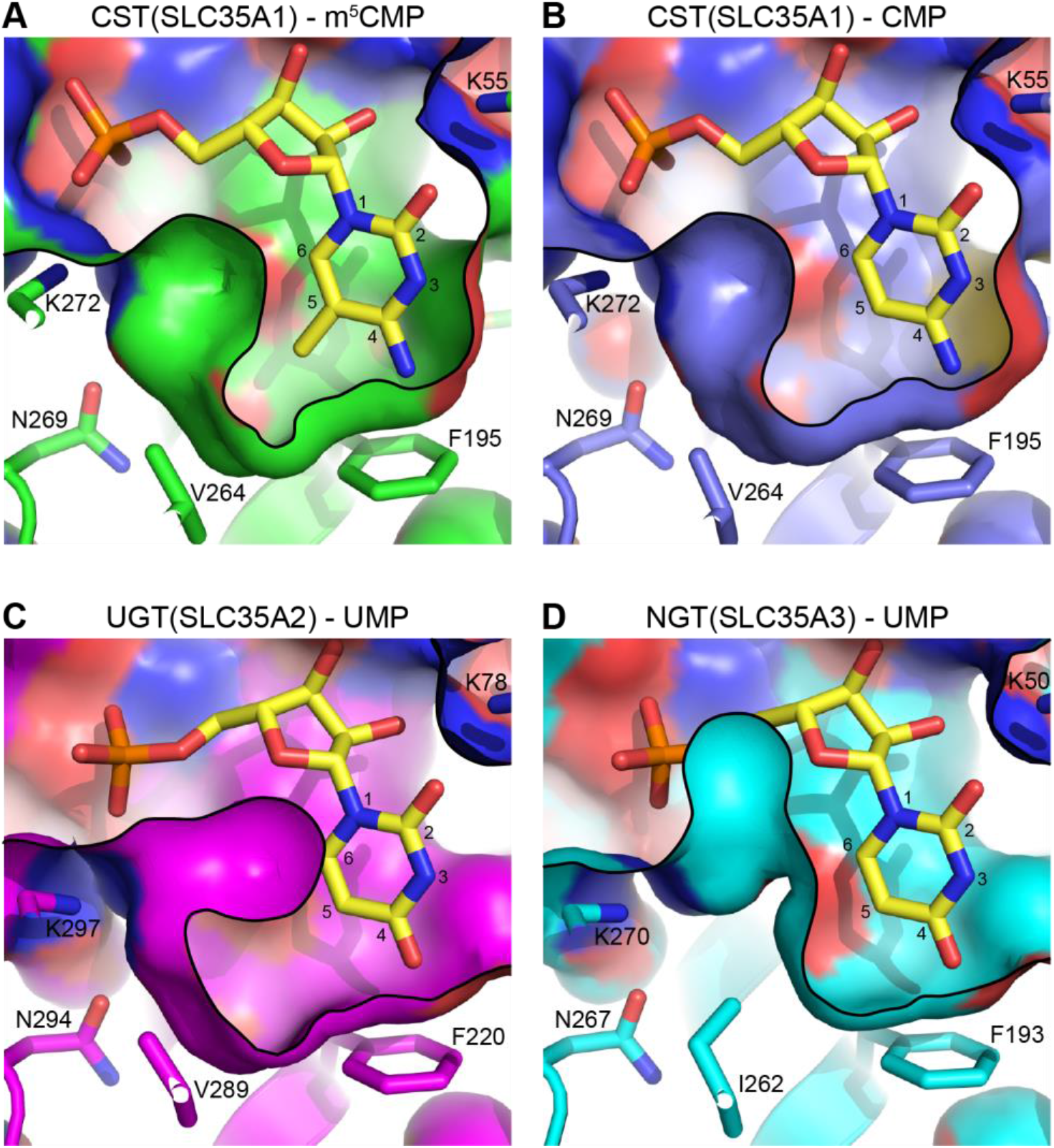
Modeling of the homologous hydrophobic pocket in UGT and NGT. **A and B)** The depiction of the hydrophobic pocket in CST is reproduced from Figure 5 for comparison. **C and D)** Structural homology models of UGT (SLC35A2, panel C) and NGT (SLC35A3, panel D) are shown with UMP docked in their substrate binding sites. The models were superimposed on the CST-m^5^CMP structure and the same view of a slice through a surface representation of the substrate-binding cavity is shown. In all panels, key residues are labeled and the atoms for the pyrimidine ring are numbered.

